# ATF4 regulates mitochondrial dysfunction and mitophagy, contributing to corneal endothelial apoptosis in Fuchs’ dystrophy

**DOI:** 10.1101/2024.11.14.623646

**Authors:** Saba Qureshi, Stefan Y. Kim, Stephanie Lee, Lukas Ritzer, William Steidl, Gayle Joliet Krest, Anisha Kasi, Varun Kumar

## Abstract

**PURPOSE:** Endoplasmic reticulum (ER) stress, mitochondrial dysfunction, and mitophagy are known to contribute independently to corneal endothelial cell (CEnC) apoptosis in Fuchs’ endothelial corneal dystrophy (FECD). However, the role of a specific ER stress pathway (PERK-ATF4-CHOP) in regulating these events is unknown. This study aims to investigate the role of ATF4 in regulating mitochondrial dysfunction and mitophagy, which ultimately leads to CEnC apoptosis in FECD.

**METHODS:** Human corneal endothelial cell line (21T), Fuchs’ corneal endothelial cell line (F35T), and primary human corneal endothelial cells were treated with ER stressor tunicamycin (Tun). ATF4 siRNA was used to knock down ATF4 in 21T cell line and primary corneal endothelial cells. Mitophagy and apoptotic proteins were analyzed using Western blotting. *ATF4^+/-^* and *ATF4 ^+/+^* mice were irradiated with UVA to assess ER stress and corneal endothelial apoptosis.

**RESULTS:** F35T cell line had significantly increased expression of the ER stress pathway as well as caspase-mediated apoptotic molecules compared to 21T at baseline, which further increased after tunicamycin treatment. F35T cells exhibited significantly decreased ATP and MMP, and increased mitochondrial fragmentation, which was further exacerbated after Tunicamycin. F35T cell line also demonstrated inhibition of mitophagy, similar to 21T, after treatment with Tunicamycin, despite the upregulation of mitophagy initiators. ATF4 knockdown attenuated ER and mitochondrial stress proteins, rescued mitochondrial membrane potential (MMP) loss, downregulated mitochondrial fragmentation, activated mitophagy, and prevented cell death under chronic ER stress. *ATF4^+/-^*mice had increased CE numbers, with improved cellular morphology and decreased ER stress CHOP expression, compared to ATF4+/+ mice post-UVA.

**CONCLUSIONS:** Pro-apoptotic ATF4 induction following ER stress disrupts mitochondrial function, leading to mitophagy inhibition and CEnC apoptosis. This study highlights the importance of ATF4 in ER-mitochondrial crosstalk and its contribution to CEnC apoptosis in FECD.

## INTRODUCTION

Fuchs’ endothelial corneal dystrophy (FECD) is a heritable degenerative disorder of the corneal endothelium (CE) that affects about 4% of people over 40 and has a higher incidence in females.^1^ The corneal endothelium consists of a single layer of hexagonal cells that line the back surface of the cornea. These cells are essential for maintaining corneal transparency by continuously pumping ions. Proteomic analysis has demonstrated that in FECD, extracellular membrane (ECM) dysregulation leads to aberrant organization of the corneal endothelial layer and Descemet’s membrane.^2^ The progressive loss of CEnCs, primarily due to apoptosis, is associated with alterations in cellular junctional complexes and the accumulation of extracellular matrix protrusions in Descemet’s membrane (DM), known as guttae, which are the key feature of FECD. The molecular pathogenesis of FECD is an area of active research. FECD is the leading cause of corneal transplantation worldwide, with no available pharmacological treatments. Thus, there is an unmet need to understand the disease pathogenesis in order to develop effective pharmacological interventions. Prior work indicates that mitochondrial dysfunction contributes to FECD development.^3,4^ Other research has emphasized the importance of ER stress in promoting the pathogenesis of FECD.^5,6^ While ER-mitochondrial crosstalk has been shown to play a role in the development of neurodegenerative diseases such as Parkinson’s disease, Huntington’s disease, amyotrophic lateral sclerosis, and Alzheimer’s disease, among others^7–10^, there has been significantly less research regarding the role of ER-mitochondrial crosstalk in the development of FECD.

Our lab previously demonstrated that ER stress leads to mitochondrial dysfunction and contributes to CEnC apoptosis.^11^ Specifically, our lab showed that induction of ER stress in 21T cell line via tunicamycin led to upregulation of three ER stress pathways named PERK-eIF2ɑ-ATF4-CHOP, ATF6, and IRE1ɑ-XBP1^12^ along with increased apoptotic proteins. We confirmed that this ER stress induction in the 21T cell line led to decreased ATP production, loss of MMP, and increased mitochondrial fragmentation. While our research showed upregulation of the well-studied PERK-eIF2ɑ-ATF4-CHOP pathway, further research needs to be performed to elucidate the specific role of ATF4 (a major regulatory transcription factor of the PERK-CHOP pathway) in regulating ER-mitochondrial crosstalk and its contribution to FECD pathophysiology.

ATF4 is a transcription factor that contributes to the integrated stress response (ISR), which allows eukaryotic cells to maintain homeostasis in response to cellular stresses.^13,14^ ATF4 plays a role in cellular homeostatic maintenance and organellar signaling, which may have implications for the mechanistic role of ATF4 in ER-mitochondrial crosstalk and the development of FECD, and warrants further investigation. Specifically, ATF4 is upregulated in response to crosstalk between the ER and mitochondria, as research has shown that mitochondrial dysfunction can promote ER stress and, thus, upregulation of the unfolded protein response (UPR), which affects downstream transcription.^15^ ATF4 is also a key player in the mitochondrial stress response, as it upregulates cytoprotective genes upon induction of mitochondrial stress.^16^ ER-mitochondrial crosstalk has been shown to upregulate mitophagy upon induction of ATF4 by toxic hexavalent chromium.^17^ Apart from mitophagy, ATF4 facilitates autophagy in the setting of ER stress by transcriptionally regulating proteins that contribute to autophagosome formation.^18^ For instance, ATF4 promotes and regulates autophagy in mammalian cells in response to tunicamycin-induced ER stress.^19^ In addition to its role in autophagy, ATF4 plays a role in apoptosis by enabling the degradation of the X-linked inhibitor of apoptosis protein (XIAP) during chronic ER stress, thus promoting cell death.^20^ Research has demonstrated that ultraviolet (UV) radiation may also play a role in the development of FECD. ^21,22^ UV radiation can also lead to decreased ATF4 expression, a finding that warrants further investigation.

While prior research establishes the role of ATF4 in organellar crosstalk and cellular homeostasis, there is a need for research that examines the mechanistic contribution of ATF4 to FECD pathogenesis. In our lab’s previous publication, we discussed the crosstalk between ER and mitochondria in the corneal endothelium. In this paper, we hypothesize that ATF4 plays a crucial role in ER-mitochondrial crosstalk, particularly in mitochondrial stress and mitophagy, which ultimately leads to apoptosis in FECD. This is the first study to demonstrate a differential effect on mitochondrial dysfunction and mitophagy after ER stress induction on normal and Fuchs’ corneal endothelial cell lines. This study specifically investigates the role of ATF4 in mitochondrial stress and apoptosis and Parkin-mediated mitophagy. Finally, this study examines the effect of genetic loss of ATF4 on CE cell viability after an Ultraviolet A (UVA)-induced FECD model *in vivo*.

## METHODS

### Normal and Fuchs’ cell line culture

Human corneal endothelial cell line (21T) (a kind gift from Ula V. Jurkunas, Harvard University) was maintained in Chen’s medium containing Opti-MEM (cat no. 51985034; Thermo Fisher, Waltham, MA, USA), 8% fetal bovine serum (cat no. A3840002; Thermo Fisher), 0.1 gm calcium chloride (cat no. C7902; Sigma-Aldrich, St. Louis, MO, USA), 0.4 gm chondroitin sulfate (cat no. C9819, Sigma Aldrich), gentamycin (0.5%) (cat no. 15750078; Thermo Fisher) antibiotic-antimycotic (1%) (cat no. 15240096; Millipore Sigma, Burlington, MA, USA), 25 μL of epidermal growth factor (cat no. GF144; Thermo Fisher) 50 μg of bovine pituitary extract (cat no. 50-753-3049; Fisher Scientific). Cells were grown on a T75 flask (cat no. 156499; Thermo Fisher) coated with FNC coating mix (cat no. 0407; Athena Enzyme Systems, Baltimore, MD, USA) passaged every two or three days and maintained in an incubator containing 5% CO2 at 37°C. The culture of Fuchs’ corneal endothelial dystrophy cell line 35 (F35T, 1500 CUG repeats) (a generous gift from Dr. Albert S. Jun, Johns Hopkins University) was maintained in Opti-MEM (cat no. 51985034; Thermo Fisher) supplemented with 8% fetal bovine serum, 0.1 gm calcium chloride (cat no. C7902; Sigma-Aldrich), 0.4 gm chondroitin sulfate (cat no. C9819; Sigma-Aldrich), gentamycin (0.5%) (cat no. 15750078; Thermo Fisher), antibiotic-antimycotic (1%) (cat no. 15240096; Millipore Sigma), 25 μL of epidermal growth factor (cat no. GF144; Thermo Fisher), 100 μL of nerve growth factor (cat no. 13257-019; Thermo Fisher), and 10 mg of ascorbic acid (cat no. 255564; Sigma-Aldrich). Cells were grown on a T25 flask (cat no. 156340; Thermo Fisher) coated with FNC coating mix (cat no. 0407; Athena Enzyme Systems), passaged every two or three days, and maintained in an incubator containing 5% CO_2_ at 37°C.

### Human primary corneal endothelial culture

We isolated corneal endothelial cells from human donor corneal tissues (Eversight, Chicago, IL, USA) and established primary cell culture. Corneas were washed with M199 medium (Gibco) supplemented with gentamycin (0.5%) and antibiotic-antimycotic (1%). The corneal endothelium layer was peeled off from Descemet’s membrane (DM), stripped into smaller pieces, and placed in a tube with fetal bovine serum (15%), antibiotic-antimycotic (1%), gentamycin (0.5%) and it was incubated for 2-5 days. It was then centrifuged at 3000 rpm for 6 min at RT, the supernatant was discarded, and the cells/DM strips were rinsed with HBSS by gentle flicking, re-pelleted, added 0.02% EDTA, and incubated at 37°C for 1 h. Then the tissue/cells were passed through a glass Pasteur pipette 15-20 times gently and added 2 ml of growth medium and centrifuged at 3000 rpm for 6 min at RT, supernatant was discarded, and pellet was resuspended in 500 μL of growth medium. The primary cell culture was maintained in growth medium constituted of Opti-MEM-1 (Gibco) supplemented with 8% FBS, 5 ng/ml human recombinant EGF (Peprotech), 20 ng/ml human recombinant NGF (Peprotech), 100 μg/ml bovine pituitary extract (Biomedical tech), 0.5 mM Ascorbic acid 2-phosphate (Sigma), 200 mg/L CaCl_2_, 0.08% chondroitin sulphate, 0.5% gentamycin, 1% antibiotic-antimycotic, passaged every 2 days and maintained in an incubator containing 5% CO_2_ at 37°C.

### siRNA transfection

21T, F35T, and primary corneal endothelial cells were transfected with silencer-select negative control siRNA (cat no. 4390846) or ATF4 siRNA (cat no. 4392420), (both Thermo Fisher Scientific, CA, USA) by using a Lipofectamine RNAi Max transfection reagent (Invitrogen, Life Technologies, CA, USA) according to manufacturer’s protocol. At 48 h after transfection, the cells were subjected to different doses of tunicamycin based on the needs of the different experiments.

### MTT assay for cell viability

The primary corneal endothelial cells (1 × 10^4^ cells/well) were seeded and grown in 96-well plates (cat no. 701003; Fisher Scientific). Cells were transfected with ATF4 siRNA and then treated with dimethyl sulfoxide (DMSO) (0.2%) and tunicamycin (10 μg/mL) and incubated for 24 hours. We performed MTT (3-(4,5-dimethylthiazol-2-yl)-2-5-diphenyltetrazolium bromide) assay (cat no. V13154; Thermo Fisher) for determining cell viability after different tunicamycin treatments as per the manufacturer’s protocol.

### Measurement of MMP

MMP was measured using the TMRE-Mitochondrial Membrane Potential Assay Kit (ab113852; Abcam) as per the manufacturer’s protocol. 21T and F35T (1 × 10^4^ cells/well) cells were seeded in 96-black well flat bottom plates (cat no. 3916; Corning, Inc., Corning, NY, USA) until 70% to 80% confluency, then treated with and without different concentrations of tunicamycin (0.01, 0.1, 1 and 10 μg/mL) and then incubated for 48 hours. Then, fluorescence (ex/em: 549/575 nm) was measured using a plate reader (Varioskan LUX; Thermo Fisher). We also transfected primary corneal endothelial cells with ATF4 siRNA for ATF4 knockdown and assessed MMP.

### Measurement of Complex V (ATP Synthase) activity for ATP production

21T cells were seeded in 150 mm plates until 70-80% confluent, and corresponding treatments were applied to the cells for 48 hours. Post-treatment, cells were pelleted, and sample preparation, along with measurement of Complex V activity, was performed using the ATP Synthase Enzyme Activity Microplate Assay Kit as per manufacturer’s protocol (cat no. ab109714; Abcam, Cambridge, UK). Absorbance was measured at OD 340 nm at 30°C using a microplate reader (Varioskan LUX; Thermo Fisher).

### Caspase 3/7 assay

21T and F35 cells (1 × 10^4 cells/well) were seeded in white walled 96-well plates, until 70% confluency. The cells were then treated with DMSO (0.2%) or tunicamycin (10 μg/ml) and incubated for 24 hours. After incubation, cells were treated with Caspase-Glo 3/7 reagent (G8090, Promega) according to manufacturer protocol. Then the luminescence was measured using a plate reader (Varioskan LUX; Thermo Fisher).

### Transmission Electron Microscopy

21T and F35T (4 × 10^4^ cells/well) were seeded in 8 well Permanox Lab-Tek Chamber Slides (cat no. 70413; Fisher Scientific), grown to 80% confluency, and treated with 10 μg/mL of tunicamycin. For the transfection experiment, HCEnC-21T was transfected with negative control siRNA or ATF4 siRNA and then treated with 10 μg/mL of tunicamycin. Cells were fixed with 2% paraformaldehyde/2.5% glutaraldehyde in 0.1M sodium cacodylate solution (cat no. 15960-01, Electron Microscopy Sciences (EMS); Fisher Scientific) at 4°C, rinsed in 0.1 M sodium cacodylate buffer, then post-fixed with 2% osmium tetroxide/1.5% potassium ferricyanide in 0.1 M sodium cacodylate, and en bloc stained with 2% uranyl acetate in dH2O. Cells were dehydrated in an ethanol series (25% EtOH/dH2O up to 100% EtOH), infiltrated through an ascending ethanol/resin series (cat no. 14120, EMS Embed 812 Kit; Fisher Scientific) and placed in pure resin overnight. Chambers were separated from the slides, and a modified BEEM embedding capsule (EMS, cat no. 70000-B; Fisher Scientific) was placed over defined areas containing cells. Capsules were filled with a drop of pure resin and placed in a vacuum oven to polymerize at 60°C for several hours. Epon resin was added to fill the capsules and polymerized for 72 hours in the vacuum oven. Immediately after polymerization, capsules were snapped from the substrate with pliers to dislodge the cells from the slide. Semithin sections (0.5 μm) were obtained using a Leica UC7 ultramicrotome (Leica), counterstained with 1% toluidine Blue, placed under a coverslip, and viewed under a light microscope to identify successful dislodging of cells. Ultra-thin sections were collected on copper 300 mesh grids using a Coat-Quick adhesive pen. Sections were counter-stained with 1% uranyl acetate and lead citrate and imaged on an HT7500 transmission electron microscope (Hitachi HighTechnologies, Tokyo, Japan) using an AMT NanoSprint12 12-megapixel CMOS transmission electron microscopy (TEM) Camera (Advanced Microscopy Techniques, Danvers, MA, USA). Final images (20–30 images per condition) were collected, and the brightness, contrast, and size of these images were adjusted using Adobe Photoshop CS4 software version CS4 11.0.1 (Adobe, Inc., San Jose, CA, USA).

### Parkin plasmid transfection

21T cells were transfected with 0.4 μg of YFP-Parkin using Lipofectamine RNAi Max transfection reagent (Invitrogen, Life Technologies, CA, USA) and incubated for 30 hours. Then, we added DMSO and tunicamycin (20 μg/mL) for 6 hours. The cells were then fixed in PFA, permeabilized in PBS and Triton X, and blocked for 1 hour (normal goat serum+ 0.1 g BSA+ Tween 20). Then we incubated with mouse anti-Cytochrome C (cat no.12963S, 1:1000; Cell Signaling) overnight at 4°C. The next day, the cells were incubated with goat anti-mouse secondary antibody (Alexa Fluor 647, cat no. 1510115 1:1000; Abcam) for 1 hour. Cells were washed with PBS, stained with DAPI, and visualized on a confocal microscope.

### Immunostaining

21T (4 × 10^4^ cells/well) and F35T were seeded in 2, 4 or 8 well chamber slides to grow up to 70% to 80% confluency. For ATF4/Mitotracker staining, the cells were incubated with MitoTracker Deep Red FM (50 nM) (cat no. M22426; Invitrogen) for 30 minutes, then fixed with paraformaldehyde (PFA) (4%) (cat no. J19943.K2; Thermo Fisher) for 20 minutes, blocked for one hour in blocking buffer (5% normal goat serum + 0.3% Triton X-100 [cat no. X100; Millipore Sigma]) and then incubated with rabbit anti-ATF4 antibody (cat no. 11815S, 1:1000; Cell Signaling, Danvers, MA, USA) overnight at 4°C. The next day, the cells were washed with ice-cold phosphate buffer saline solution (PBS) (cat no. 10010072; Thermo Fisher) three times and incubated with goat anti-rabbit secondary antibody (Alexa Fluor 488, cat no. ab1550077, 1:1000; Abcam, Cambridge, UK) for 1 hour. Cells were washed again with cold-ice PBS and stained nuclei with DAPI (cat no. D1306; Invitrogen). Slides were visualized on a confocal microscope (Leica STED 3X; Leica, Wetzlar, Germany). We also used primary corneal endothelial cells to probe for cytochrome C and ATF4. A similar immunocytochemistry method was used for staining mitochondria by mouse anti-cytochrome C (cat no. 12963S, 1:1000; Cell Signaling) and co-incubated with rabbit anti-ATF4 antibody except for treatment (0.4% DMSO; 10 μg/mL Tun, 24 hours) prior to fixation. The next day, the cells were co-incubated with goat anti-rabbit secondary antibody (Alexa Fluor 488, cat no. ab1550077) and goat anti-mouse secondary antibody (Alexa Fluor 647, cat no. ab1510115, 1:1000; Abcam) for 1 hour. Cells were washed with PBS and stained with DAPI and visualized on a confocal microscope. For transfection experiment, primary corneal endothelial cells were transfected with negative control siRNA or ATF4 siRNA. At 48 h after transfection, the cells were treated with 10 μg/mL of tunicamycin for 24 hours, then followed the same staining procedure for ATF4 and cytochrome C. For the human corneal tissue immunostaining, Descemet’s membrane containing endothelial cells was stripped from human donor corneas and treated with DMSO (0.2%) and tunicamycin (10 μg/mL) and incubated for 24 hours. After treatment, fixation, and blocking, the tissues were then incubated with mouse anti-CHOP antibody (cat no. 2895S, 1:1000; Cell Signaling) and rabbit anti-ATF4 antibody overnight at 4°C. For quantifying ATF4^+^DAPI^+^ cells per visual field (×40 lens), 10 to 15 images were taken under each condition, and the total number of ATF4^+^ DAPI^+^ cells (25-30) was counted and averaged. A similar procedure was followed for calculating the percentage of fragmented mitochondria using cytochrome C. Quantification was masked to reduce the experimental bias. For ZO-1 staining, a dissected mouse cornea cup was fixed with 70% ethanol for 30 min at room temperature, permeabilized with 0.2% Triton X-100 in PBS for 10 min, and then blocked with 2% bovine serum albumin (BSA)-PBS for 15 to 30 min. The cornea cup was then incubated with the mouse anti-CHOP antibody (1:400, Cell Signaling) for CHOP immunostaining assay or an anti-ZO-1 antibody (cat no. ZO1-1A12, Invitrogen) for ZO-1 assay in 4% BSA-PBS at 4 °C overnight. The tissues were washed with PBS the following day and incubated with goat anti-mouse secondary antibody for 1 hour. Slides were made and imaged with confocal microscopy. In vivo CE quantification was performed using ImageJ software with the Cell Counter function to count stained cells within each visual frame. For each experimental condition, five images were analyzed, and cell counts from these images were averaged to obtain a representative cell count.

### Western blot

21T and F35T were treated either with or without DMSO or tunicamycin (DMSO: 0.0002, 0.002, 0.02, 0.2%; tunicamycin: 0.01, 0.1, 1, 10 μg/mL) for 24 hours for analyzing ER and mitochondrial stress proteins. Primary corneal endothelial cells were transfected with ATF4 siRNA and treated with DMSO or tunicamycin (10 μg/mL) for 24 hours for analyzing autophagy markers. Cells were lysed using RIPA lysis and extraction buffer (cat no. 89900; Thermo Fisher) containing protease and phosphatase inhibitor cocktail (cat no. 78440, 1:1000; Thermo Fisher) at 24 hours and spun in a centrifuge at 10,000g for 10 minutes at 4°C. The supernatant was collected and assessed to quantify protein concentration (A53227; Thermo Fisher). For the transfection experiment, primary corneal endothelial cells were transfected with negative control siRNA or ATF4 siRNA. At 48 h after transfection, the cells were treated with 10 μg/mL of tunicamycin. Protein 30 μg mixed with appropriate LDS sample buffer (cat no. NP0007; Thermo Fisher) and reducing agent (cat no. NP0009; Thermo Fisher) was subjected to 4% to 12% sodium dodecyl sulfate-polyacrylamide gel electrophoresis (cat no. NP032; Thermo Fisher), followed by transfer to polyvinylidene difluoride membrane (cat no. 88518; Thermo Fisher) at 30 V for 90 minutes. The blots were blocked in blocking buffer (25 mM Tris, 0.15 M NaCl, 0.05% Tween 20, 5% skim milk) for one hour at room temperature and then treated with primary antibody (p-elF2α cat no. 3597, eIF2α cat no. 5324, CHOP cat no. 2895, ATF4 cat no. 11815, GAPDH cat no. 97166, Bcl2 cat no. 4223, cleaved caspase 9 cat no. 20750, cleaved caspase 3 cat no. 9661, PARP cat no. 9532, Mfn2 cat no. 11925, PINK1 cat no. 6946, Parkin cat no. 4211, LC3A/B cat no. 12741, Tim23 cat no. 67535, LC3B cat no. 3868, VDAC cat no. 4661, P-mTOR cat no. 5536, mTOR cat no. 2983, P-Akt cat no. 4060, Akt cat no. 4691, P-AMPKα cat no. 2535, AMPKα cat no. 5831, Actin cat no. 3700; all from Cell Signaling; except Tim23 from Proteintech) overnight at 4°C. The next day, the blots were washed three to five times in TBS-Tween 20 (25 mM Tris, 0.15 M NaCl, 0.05% Tween 20) (cat no. 28360; Thermo Fisher) and HRP-conjugated secondary antibody (anti-mouse IgG cat no. 7076S, Anti-rabbit IgG cat no. 7074S; Cell Signaling) was added for one hour with similar subsequent washing. The immunoreactive bands were visualized using a Femto (cat no. 34094; Thermo Fisher) or pico chemiluminescent substrate (cat no. 34579; Thermo Fisher). The band intensity was measured and quantified using ImageJ software.

### Animals

*ATF4^+/+^, ATF4^+/-^,* mice (JAX:013072) were generated at Jackson Laboratory (Farmington, CT, USA) by embryo reconstitution and crossing with healthy wild-type-C57BL6/N mice. Monogamous pairs consisting of one male and one female ATF4+/+ and ATF4+/-mouse were crossed as per a homozygous X heterozygous breeding scheme. Litters were genotyped by TransnetYX (Cordova, TN) to identify *ATF4^+/+^* and *ATF4^+/-^* mice. Mice were housed at the Animal Housing Facility, Eye and Vision Research Institute (EVRI), Mount Sinai, New York, USA, in a controlled environment with a constant temperature, a 12-h light/dark cycle, and food and water available ad libitum. Mice were anesthetized with a combined dose of ketamine (100 mg/kg) and xylazine (20 mg/kg) administered intraperitoneally (IP). Animal studies were in accordance with the ARVO Statement for the Use of Animals in Ophthalmic and Visual Research as well as the NIH Guide for the Care and Use of Animals and were performed at EVRI with approval from EVRI Institutional Animal Care and Use Committees IACUC.

### UVA irradiation

The methods of UVA irradiation of mouse cornea were described in our earlier publication. Briefly, we created a UVA LED light source (M365LP1; Thorlabs, USA) with an emission peak of 365 nm light, 8 nm bandwidth (FWHM). The irradiance of 398 mW/cm^2^ was focused down to a 4 mm diameter illumination spot onto the mouse cornea. The time of UVA exposure was adjusted to deliver the appropriate fluence (23 minutes for 200 J/cm^2^) as measured with a thermal power sensor head (S425C, Thorlabs, USA) and energy meter console (PM100D, Thorlabs). We irradiated the right eye (OD) with UVA, while the contralateral eye (OS) served as an untreated control and was covered with retention drapes (SpaceDrapes, Inc., USA). Mouse eyes were enucleated in sterile PBS, and corneas containing Descemet’s membrane were isolated and immunostained based on experiments. We have considered sacrificing mice on day 1 and week 4 post-UVA for counting CEnC numbers and ER stress marker protein, CHOP, respectively.

## RESULTS

### Differential activation of ER stress pathway, PERK-eIF2ɑ-ATF4-CHOP, after chronic ER stress in 21T and F35T cell lines, primary CE culture, and human corneal tissues

Previous studies have observed morphological changes in the ER along with increased expression of eIF2 and CHOP proteins of PERK-eIF2ɑ-ATF4-CHOP ER stress pathway within the corneal endothelium of FECD patients.^23^ Thus, we first investigated whether there is differential activation of ER stress-related proteins in the PERK-eIF2α-ATF4-CHOP pathway, specifically eIF2α, p-eIF2α, ATF4, and CHOP proteins, in F35T and 21T cell lines after Tun-induced chronic ER stress. Western blotting data showed increased expression of peIF2α, ATF4, and CHOP in F35T compared to the 21T after Tunicamycin (Tun). **(Fig.1A, B)**. Our primary focus was on pro-apoptotic ATF4 and CHOP proteins in this study. Immunostaining data showed a significant induction of ATF4 (mainly in the nucleus, with some induction in mitochondria) in F35T compared to 21T cell line under normal physiological conditions **(Fig. 1C, D).** This is confirmed by our previous publication, in which we observed baseline activation of the PERK-EIF2α-ATF4-CHOP pathway proteins in the F35T compared to the 21T.^12^ We also investigated this by using primary endothelial cells derived from human corneal tissues. We observed increased ATF4 expression after Tun treatment compared to DMSO in primary endothelial cells **(Fig. 1E)**. To further validate these data, we used corneal tissues and investigated ATF4 and CHOP expression after Tun treatment. We demonstrated increased expression of ATF4 and CHOP proteins after treatment with Tun compared to DMSO in human corneal tissue (specifically CEnCs) using immunohistochemistry **(Fig. 1F)**. These data suggest the potential involvement of the PERK-eIF2α-CHOP ER stress pathway in the pathogenesis of FECD.

**Figure 1.**
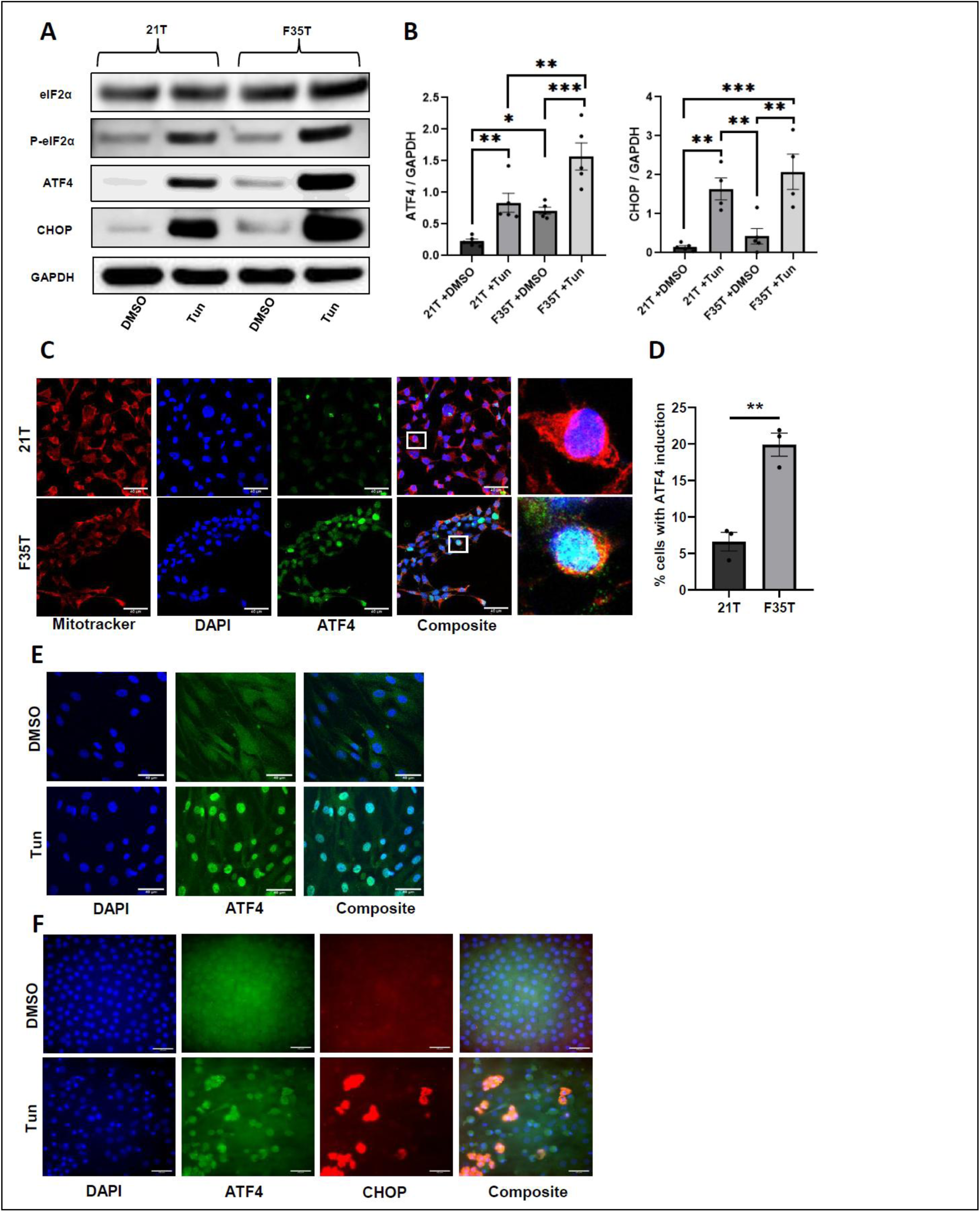
Differential activation of ER stress markers after chronic ER stress in normal human corneal endothelium cell line (21T), Fuchs (F35T) cell line, primary human corneal endothelial cells, and human corneal tissues. **(A)** Representative Western blot data showing differential activation of ER stress-related proteins (eIF2α, p-eIF2α, CHOP, and ATF4) in 21T and F35T cell lines under DMSO (0.2%) and Tun (10 µg/mL) for 24 h; DMSO was used as solvent control. **(B**) Quantification of westen blots demonstrating increased expression of ATF4 and CHOP in F35T compared to 21T after Tun (n=3-5, **P < 0.01, ***P < 0.001,one-way ANOVA with Tukey’s multiple comparison test). **(C)** Immunostaining images ATF4 (green) and mitochondria (labeled with mitotracker red) in F35T and 21T cell lines under normal physiological conditions. **(D)** Quantification of immunostaining data showing increased % of cells with ATF4 induction in F35T compared to 21T cell line under normal physiological conditions (n = 3, **P < 0.01, unpaired t-test). **(E)** Immunostaining images showing ATF4 activation after treatment with Tun (10 µg/mL, 24h) in primary human corneal endothelial cells. **(F)** Immunostaining showing ATF4 and CHOP induction after treatment with Tun (10 µg/mL, 24 h) in human corneal endothelial tissues. (scale bar: 40 μm for all immuno images).

### Differential mitochondrial dysfunction and apoptosis induced by chronic ER stress in 21T and F35T cell lines

ER stress is known to induce mitochondrial stress and dysfunction by initiating the loss of MMP, decreased ATP production, and subsequent release of cytochrome C from the mitochondria.^12^ We first investigated whether there are any baseline differences in MMP between normal and Fuchs’ cell lines. There were no differences in MMP for 21T and F35T cell lines at baseline. However, when we induced ER stress using tun, we observed a dose-dependent decrease in MMP with Tun in F35T compared to 21T cell lines (**Fig. 2A**). We also found a significantly decreased ATP production in F35T compared to 21T cell line at a basal level which further attenuated after treatment with Tun **(Fig. 2B)**. Immunostaining data also showed increased mitochondrial fragmentation in F35T cell line compared to 21T cell line under DMSO control which further increased after Tun-induced ER stress **(Fig. 2C, D)**. These data suggest that Fuchs cells exhibit higher mitochondrial damage compared to control cells under chronic ER stress. We further investigated whether mitochondrial dysfunction leads to apoptosis. Western blot and quantification data showed upregulation of PARP, cleaved caspase 3 and 9, as well as downregulation of Bcl-2 in F35T compared to 21T cell line at baseline **(Fig. 2E, F).** Next, we performed a cleaved caspase 3/7 assay and found a significant increase in cleaved caspase 3/7 activity in F35 compared to 21T under Tun **(Fig. 2G)**. Furthermore, we also found an increased trend in cleaved caspase 3 protein expression and no significant differences in Bcl-2 between F35 and 21T under Tun (**Fig. 2H**). We also found a significant increase in cleaved PARP (**Fig. 2I**) and caspase 9 **(Fig. 2J)** in F35 compared to 21T under Tun. These data suggest increased caspase-mediated apoptosis in Fuchs compared to control under chronic ER stress, possibly contributed by mitochondrial dysfunction.

**Figure 2.**
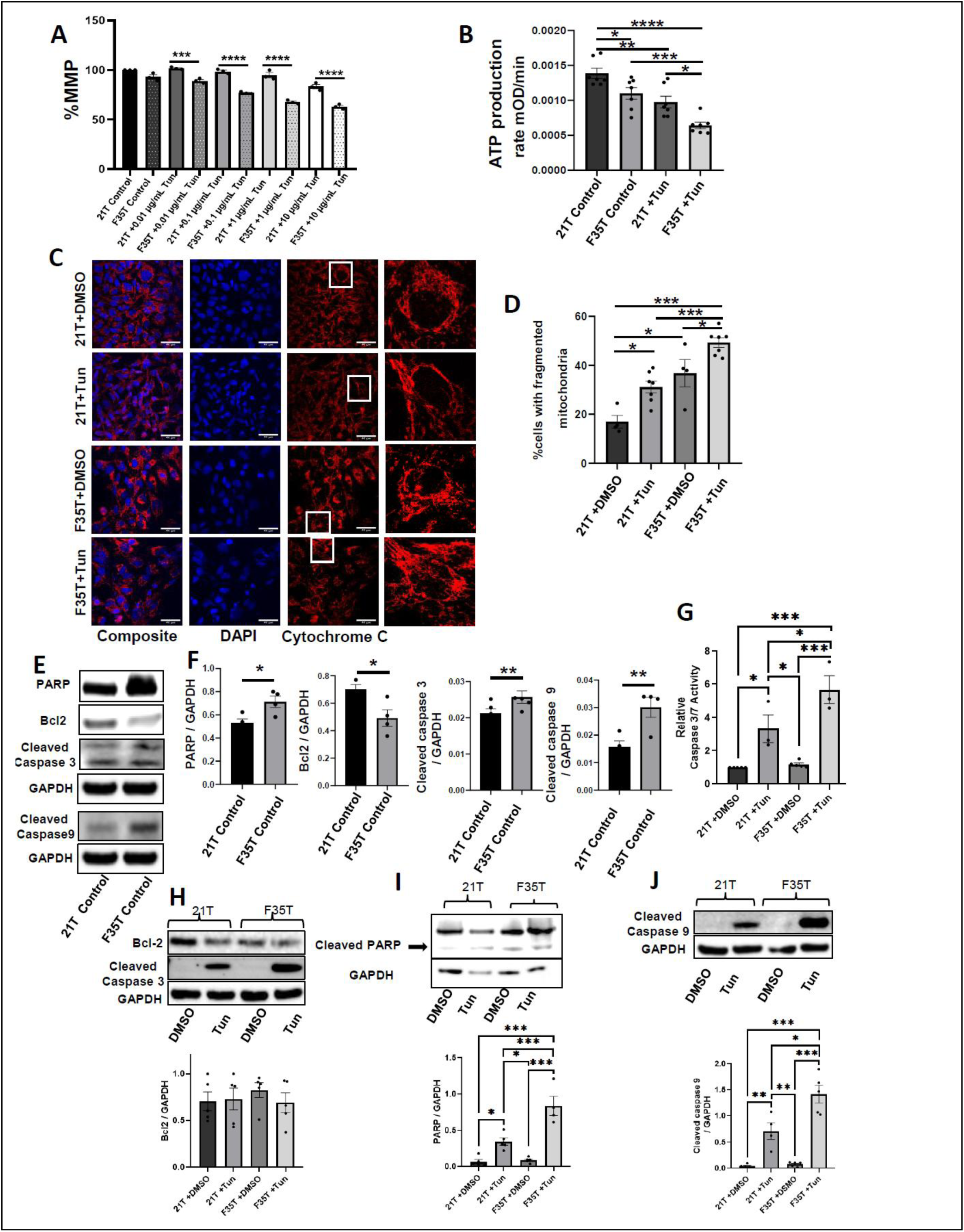
Differential mitochondrial dysfunction induced by ER stress in 21T and F35T. **(A)** Bar graph demonstrating differential MMP loss between 21T and F35T cell lines with and without Tun treatment (0.01, 0.1, 1, and 10 µg/mL Tun) (n = 3, ***P < 0.001, ****P < 0.0001, one-way ANOVA with Tukey’s multiple comparison test). **(B)** Bar graph demonstrating differences in ATP production between untreated and treated (10 µg/mL Tun) 21T and F35T cell lines; (n = 7, *P < 0.05, **P < 0.01, ***P < 0.001, ****P < 0.0001, one-way ANOVA with Tukey’s multiple comparison test). **(C, D)** Representative immunostaining and quantification showing increased mitochondrial fragmentation (observed by diffuse cytochrome C staining and loss of mitochondrial tubular networks) in F35T compared to 21T cell line after Tun treatment (10 µg/mL at 24h). F35T cell line also has significantly increased mitochondrial fragmentation compared to 21T cell under DMSO control at 24 h (n = 4, *P < 0.05, ***P < 0.001, one-way ANOVA with Tukey’s multiple comparison test, scale bar: 40 μm), **(E)** Representative Western blots showing differential activation of mitochondrial-mediated intrinsic apoptotic proteins. (Bcl2, Cleaved Cas9, Cleaved Cas3, and PARP) in 21T compared to F35T cell lines under physiological conditions. **(F)** Bar graph demonstrating increased activation of PARP, Cleaved cas 3,9 and decreased activation of anti-apoptotic protein, Bcl-2 in F35T cell line compared to 21T cell line under physiological conditions (n = 4, *P < 0.05, **P < 0.01, unpaired t-test). **(G)** Cleavage 3/7 activity assay showing significantly increased Cleaved 3/7 activity in F35T compared to 21T after Tun (10 µg/mL at 24h). **(H)** Representative western blot along with quantification showing increasing trend of cleaved caspase 3, but no change in Bcl-2 in F35T compared to 21T after DMSO (0.2%) and Tun (10 µg/mL at 24h) (n = 4, *P < 0.05, ***P < 0.001, one-way ANOVA with Tukey’s multiple comparison test). Representative western blot and quantification showing increased expression of **(I)** cleaved PARP **(J)** caspase 9 in F35T compared to 21T after DMSO (0.2%) and Tun (10 µg/mL at 24h) (n = 4, *P < 0.05, ***P < 0.001, one-way ANOVA with Tukey’s multiple comparison test, scale bar: 40 μm).

### Mitophagy activation under chronic ER stress in 21T and F35T cell lines

We hypothesized that mitochondrial dysfunction plays a crucial role in initiating mitophagy events. Thus, we determined the expression of a major mitophagy-contributing protein/mitochondrial fusion protein, mitofusin 2 (Mfn2), and mitophagy initiator proteins (PINK1, Parkin, LC3II/I) in F35T and 21T after chronic ER stress. We found a significant upregulation of Parkin and LC3, but a downregulation of PINK1 and Mfn2 in F35T compared to 21T at baseline **(Fig. 3A, B**). Mfn2 was further attenuated in both 21T and F35T cell lines after Tun **(Fig. 3C, D)**. Moreover, LC3II/I, specifically LCII denoted by the lower band **(Fig. 3C, D)** and Parkin **(Fig. 3E, F)**, significantly increased in both F35T and 21T after Tun treatment. However, there was a downregulation of PINK1 in both F35T and 21T cell lines after Tun **(Fig. G, H).** These data suggest that the initial steps in mitophagy are already activated at baseline in the Fuchs cell line, requiring no further activation of mitophagy initiator proteins (Parkin, LC3) and downregulation of the mitophagy contributor protein (Mfn2). To confirm Parkin-mediated mitophagy activation under chronic ER stress, we transfected 21T with YFP-Parkin and found a significant increase in Parkin localization in the fragmented mitochondria of 21T after Tun **(Fig. 3I, J)**. Although Parkin-mediated mitophagy initiation is reported in Fuchs. However, it is unknown in Fuchs whether the mitophagy process is completed, resulting in fewer mitophagosomes containing damaged mitochondria. This is more relevant here since in cardiac^24^ and skeletal diseases^25^, loss of Mfn2 is associated with the inhibition of mitophagy, resulting in increased mitophagosomes containing damaged mitochondria. Our TEM data suggest an increase in mitophagosomes in F35 and 21T cells after Tun treatment (**Fig. 3K),** indicating inhibition of mitophagy in both cell lines and no differences in mitophagy regulation following chronic ER stress. This contrasted with the activation of mitophagy initiators in both cell lines after chronic ER stress (**Fig. 3C-F**).

**Figure 3.**
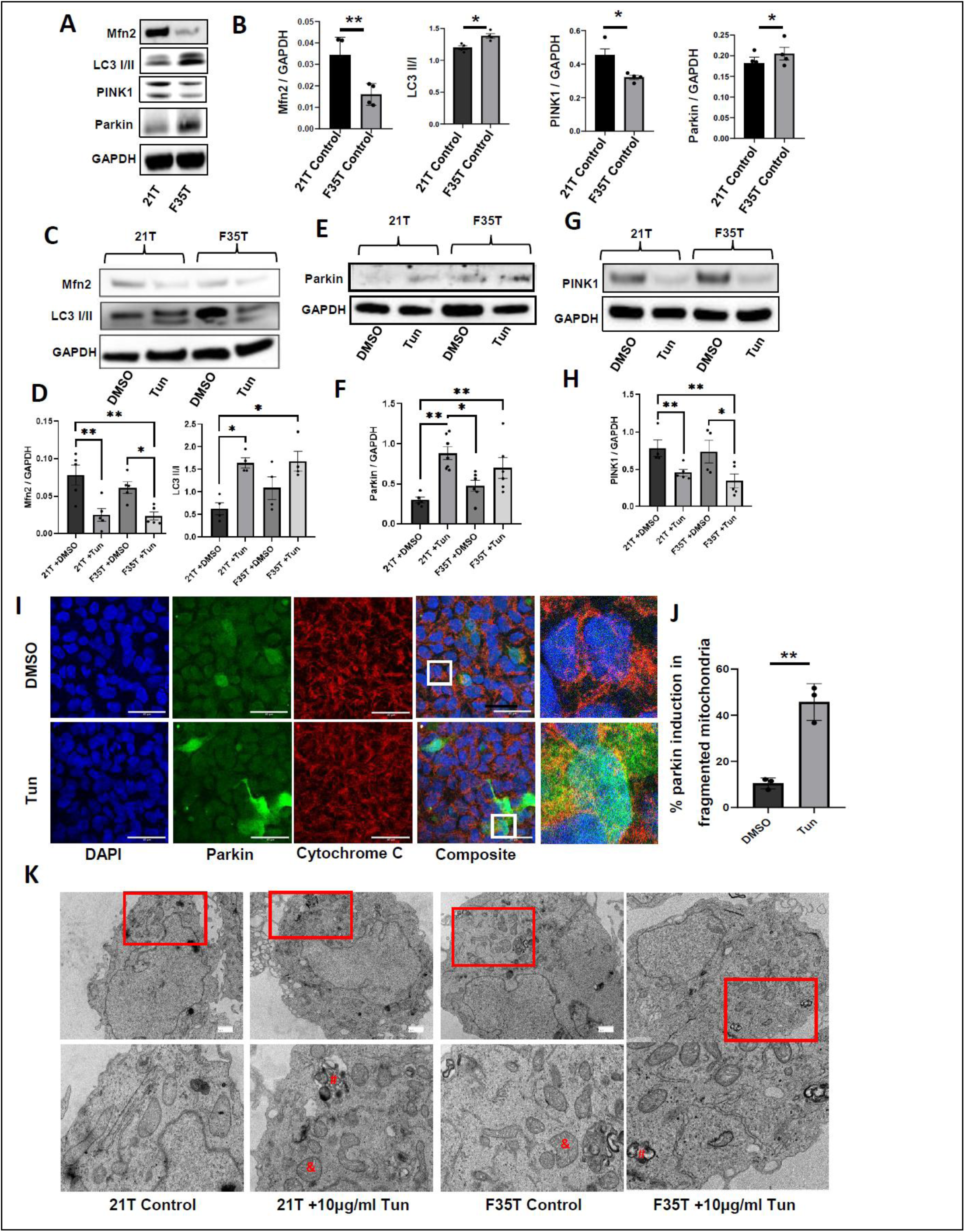
Altered mitophagy under chronic ER stress in 21T and F35T cell lines. **(A)** Representative western blots showing differential expression of Mfn2, LC3 I/II, PINK1, and Parkin in 21T and F35T cell lines under physiological conditions. **(B)** Bar graph demonstrating upregulation of parkin and LC3B, and downregulation of PINK1 and Mfn2 in F35T cell line compared to 21T cell line under normal conditions. Representative western blot and bar graph demonstrating **(C, D)** decreased expression of Mfn2 and increased expression of LC3 II/I in **(E, F)** increased expresssion of Parkin **(G,H)** decreased expresssion of PINK1 in F35T and 21T cell line under tun (10 µg/mL for 24 h). (n = 4-7, *P < 0.05, **P < 0.01, unpaired t-test for two groups and one-way ANOVA with Tukey’s multiple comparison test for more than two groups). **(I)** Immunostained images showing increased parkin activation (green) and cytochrome C release (red) after treatment with Tun (20 µg/mL for 6 h) in 21T cell line (scale bar: 40 μm). **(J)** Quantification of immunostaining data showing % of fragmented mitochondria with Parkin induction post-Tun treatment (20 µg/mL for 6 h) compared to DMSO (n = 3, **P < 0.01, unpaired t-test). **(K)** TEM images showing swollen, fragmented mitochondria (denoted by #), along with a similar number of mitophagosomes (denoted by $), in F35T and 21T cell lines under treated conditions (10 µg/mL Tun for 24 h). Red-colored insets show enlarged images below for better visualization (scale bar: 1 μm).

### ATF4 knockdown decreases pro-apoptotic ER and mitophagy mediator proteins, attenuates mitochondrial fragmentation and MMP loss, and increases cell viability under chronic ER stress

As described earlier, ATF4 is a transcription factor that is activated in response to chronic ER stress in many diseases.^26,27^ To investigate the role of ATF4 in regulating ER stress-mediated mitochondrial dysfunction, we knocked down ATF4 expression using ATF4 siRNA and found a significant knockdown of ATF4 after chronic ER stress. ATF4 is recognized for its role as a protective factor during cellular stress. Its pro-death function has been linked to its regulation of CHOP, which is involved in triggering apoptosis in response to ER stress.^28,29^ We observed decreased expression of ATF4, CHOP along with other caspases (cleaved cas 3 and 9) under ATF4 siRNA compared to control siRNA after Tun, suggesting ATF4 knockdown rescues the cell death-related molecules after chronic ER stress (**Fig. 4A**). We also focused on mitophagy contributing protein, Mfn2, a mitochondrial fusion protein, which plays a crucial role in regulating mitochondrial dynamics following stress. We also investigated another mitophagy contributing protein, Tim23, which is an integral inner membrane protein essential for mitochondrial import. ^30^ Tim23 depletion from cells results in a defect in import.^31^ Mfn2 and Tim 23 increased in ATF4 siRNA compared to control siRNA after Tun **(Fig. 4A).** This data demonstrates that ATF4 regulates mitophagy-regulating molecules, possibly contributing to apoptosis. Parkin and PINK1 are key regulators that control mitophagy and maintain mitochondrial homeostasis.^32^ PINK1/Parkin-mediated mitophagy is typically viewed in the context of mitochondrial depolarization by facilitating the turnover of damaged mitochondria. We also investigated mitophagy initiator proteins (PINK1 and Parkin) and found that PINK1 levels decreased and Parkin levels increased under ATF4 siRNA compared to control siRNA after tun treatment **(Fig. 4A).** This data suggests that ATF4 activation under chronic ER stress mediates the initiation of the Parkin-PINK1 mitophagy pathway. Mitophagy is typically initiated following the loss of MMP and mitochondrial fragmentation in response to chronic stress. Thus, we determined MMP and found that the loss of MMP upon chronic ER stress was rescued after ATF4 knockdown under Tun **(Fig. 4B).** Cell viability significantly increased under ATF4 siRNA compared to control siRNA after Tun, as shown by MTT assay **(Fig. 4C).** The immunostaining data in primary corneal endothelium cells showed that ATF4 gets induced and translocates to fragmented mitochondria after Tunicamycin (also shown in **Fig 3I,J**) and ATF4 knockdown significantly decreased fragmented mitochondria compared to control siRNA after tun **(Fig. 4D, E).** These data suggest that ATF4 regulates mitochondrial dysfunction, contributing to CEnC apoptosis under chronic ER stress.

**Figure 4.**
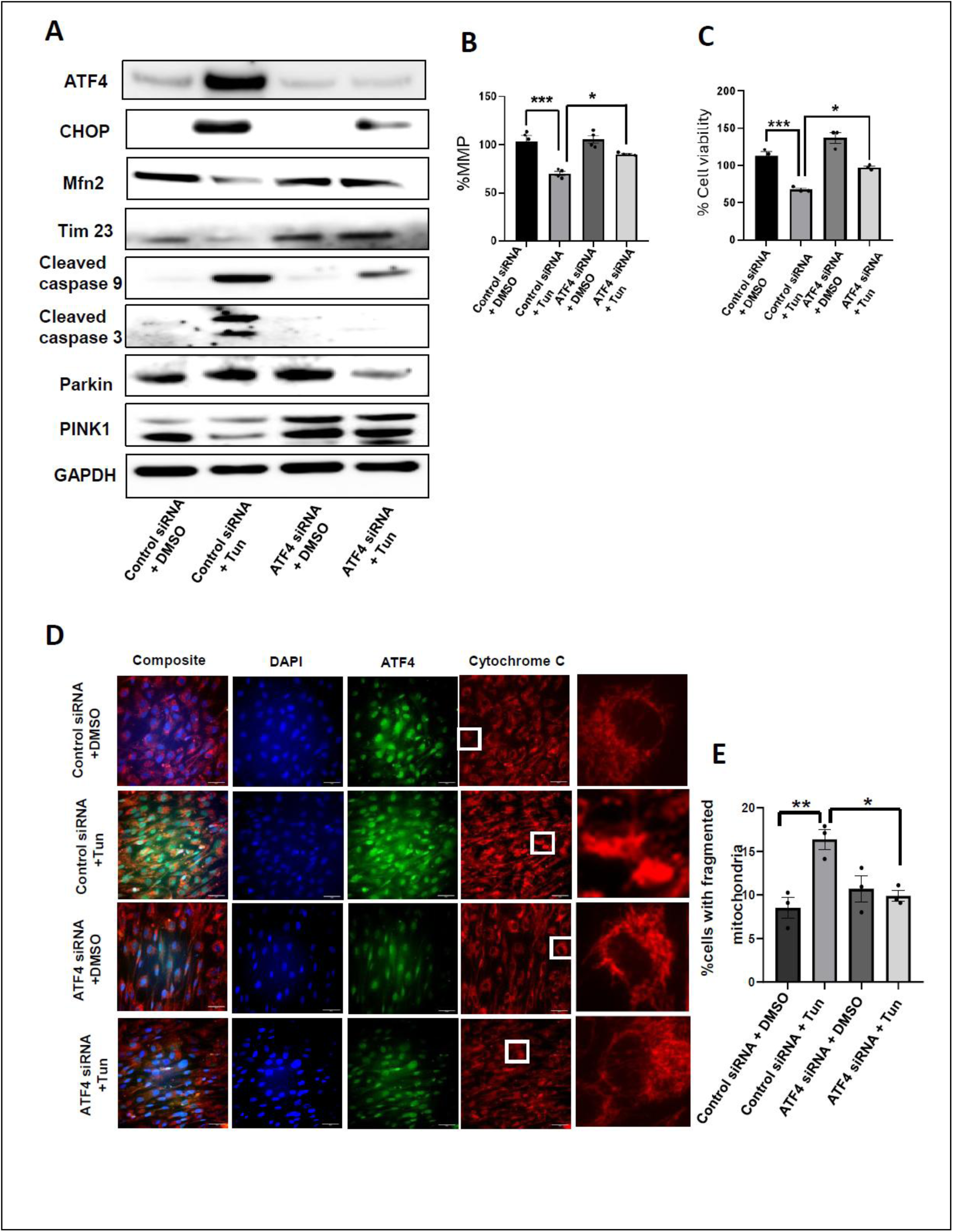
ATF4 knockdown attenuates ER stress, MMP loss, mitophagy-mediator proteins, mitochondrial fragmentation, and apoptosis under chronic ER stress. **(A)** Western blots showing upregulation of pro-apoptotic ER stress proteins (ATF4, CHOP), downregulation of mitochondria mass related proteins (Mfn2, Tim23), upregulation of apoptotic proteins (cleaved caspase 3, cleaved caspase 9) and differential regulation of mitophagy mediator proteins (increaed Parkin and decreased PINK1) in Tun treated groups (10 µg/mL, 24h) compared to DMSO (0.2%, 24 h) under control siRNA and reversal of these ER, mitochondrial, mitophagy as well apoptosis proteins expression in ATF4-knockdown groups, compared to control siRNA under Tun. **(B)** Bar graph demonstrating MMP loss post Tun treatment (10 µg/mL Tun) comapred to DMSO (0.2%) under control siRNA at 48 h in primary corneal endothelial cells while ATF4 knockdown rescues MMP loss compared to control siRNA under Tun **(C)** Bar graph demonstrating loss of cell viability using MTT post Tun treatment (10 µg/mL Tun) compared to DMSO (0.2%) under control siRNA and rescue of cell death post ATF4 siRNA treatment compared to control siRNA under Tun in primary corneal endothelial cells at 24 h; (For MMP and MTT, n = 3-4, *P < 0.05, ***P < 0.001, one-way ANOVA with Tukey’s multiple comparison test). **(D)** Immunostaining for ATF4 (green) and cytochrome c (red), DAPI (blue) in primary human corneal endothelial cells for control and ATF4 siRNA under DMSO (0.2%) and Tun (10 µg/mL) at 24 h (scale bar: 40 μm). **(E)** Quantification of immunostaining data showing significant increase in % of fragmented mitochondria post tunicamycin treatment compared to DMSO under control siRNA and rescue of mitochondrial fragmentation after ATF4 siRNA comparfed to control siRNA under Tun (10 µg/mL Tun) (n = 5, *P < 0.05, **P < 0.01, one-way ANOVA with Tukey’s multiple comparison test).

### ATF4 knockdown augments the mitophagy process despite a decrease in mitophagy-initiators under chronic ER stress, thereby contributing to CEnC viability

To determine whether mitophagy initiators are affected in the mitochondrial fraction under ATF4 manipulation after chronic ER stress, we knock down ATF4, induce chronic ER stress, isolate mitochondria from 21T cells, and probe for Parkin, PINK1, and LC3II. We found that Parkin and LC3II were upregulated, while PINK1 was downregulated, in Tun-treated cells compared to DMSO-treated cells under control siRNA, suggesting chronic ER stress mediates activation of mitophagy initiators. **(Fig. 5A).** When we knocked down ATF4, Parkin and LC3B were downregulated, suggesting inhibition of mitophagy initiators. This finding led us to suspect that mitophagy is inhibited after ATF4 knockdown under chronic ER stress, which could be detrimental to the cells, given that many diseases involve mitophagy inhibition contributing to the loss of cell viability. However, this is not observed in our data, as ATF4 knockdown increases cell viability after chronic ER stress (Fig. 4C). This suggests that even though mitophagy initiators (Parkin, LCII) are inhibited under ATF4 knockdown after chronic ER stress, however, the existing level of mitophagy initiators might be enough to complete the whole mitophagy process where we should not see accumulation of damaged mitochondria in mitophagosomes. Therefore, we performed TEM and analyzed mitophagosomes in ATF4 and control siRNA-treated 21T under DMSO and Tun conditions. Our TEM data suggests a significant increase in mitophagosomes under Tunicamycin compared to DSMO under control siRNA **(Fig. 5B, C).** This suggests that mitophagy is inhibited under chronic ER stress induced by Tun. ATF4 knockdown significantly decreases the number of mitophagosomes compared to control siRNA after tunicamycin treatment **(Fig. 5B, C),** suggesting that ATF4 induction under chronic stress inhibits mitophagy, which may contribute to the loss of cellular viability.

**Figure 5.**
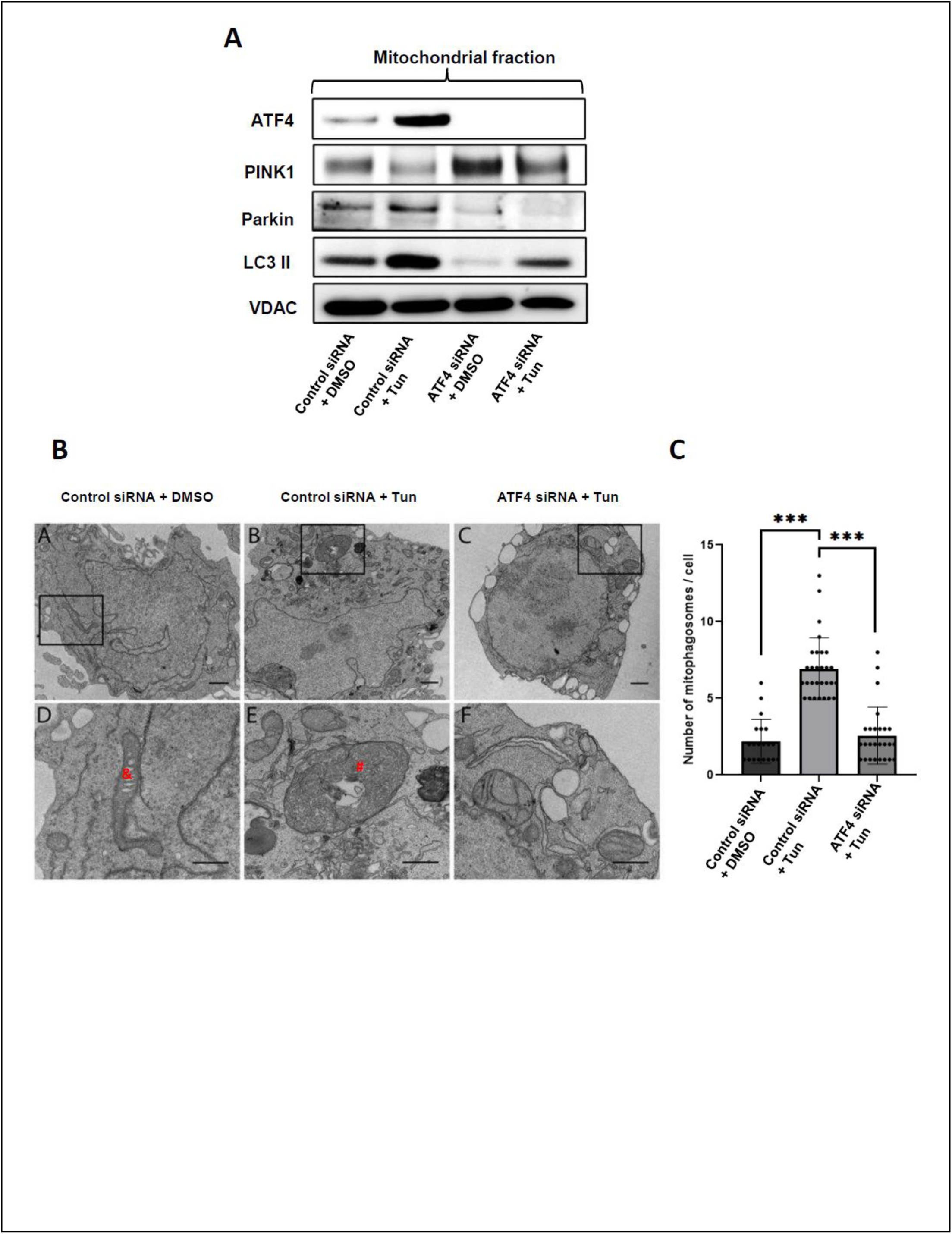
ATF4 knockdown decreases parkin-mediated mitophagy mediator proteins; however, it activates mitophagy under ER stress. **(A)** Western blot data from isolated mitochondrial fraction of 21T cells showing increased expression of ATF4, differential activation of mitophagy mediators (increased Parkin and LC3II and decreased PINK1) post ER stress (0.1 µg/mL Tun at 24 h) under control siRNA, and reversal of expression of mitophagy mediators under ATF4 siRNA compared to control siRNA post Tun. **(B)** TEM images showing mitochondria (denoted by $), mitophagosomes (denoted by #) under control siRNA and ATF4 siRNA with DMSO and Tun treatment (10 µg/mL) (C) Quantitive bar graph showing increased number of mitophagsomes in Tun condition compared to DMSO under control siRNA and a signiifcant decrease in number of mitophagosomes under ATF4 siRNA groups compared to control siRNA post Tun. (A total of 20-25 cells have been imaged using TEM for counting mitophagosomes, scale bar: 1 μm).

### ATF4^+/-^ mice have significantly decreased CHOP activation and increased corneal endothelial cell number compared to ATF4 ^+/+^ mice post-UVA

We used our established UVA-induced mouse model of Fuchs’ dystrophy^22^ to investigate the loss of corneal endothelial cells in *ATF4^+/+^* and *ATF4^+/-^* mice corneas. For this, we irradiated *ATF4 ^+/+^* and *ATF4 ^+/-^* mice’ eyes with UVA (200 J/cm^2^) and observed the change in morphology of corneal endothelial (CE) cells, as outlined by ZO-1 immunostaining, and counted the number of mouse CE cells. We found that *ATF4^+/-^* mice corneas have a significantly increased number of corneal endothelial cells with more hexagonal morphology compared to *ATF4 ^+/+^* mice post-UVA (**Fig. 6A, B**), which suggested ATF4 regulation of CE cell viability *in vivo*. To investigate whether the increased CE cell number in ATF4+/-mice also results in decreased CHOP, a pro-apoptotic ER stress molecule, we calculated the percentage of CE cells with CHOP activation. We found a significantly increased % of CE cells with CHOP in *ATF4^+/+^* compared to *ATF4^+/-^* mice at day 1 post-UVA, with no differences in non-UVA control (**Fig. 6C, D**). This suggests that ATF4 regulates CE cell viability by regulating CHOP activation *in vivo*.

**Figure 6.**
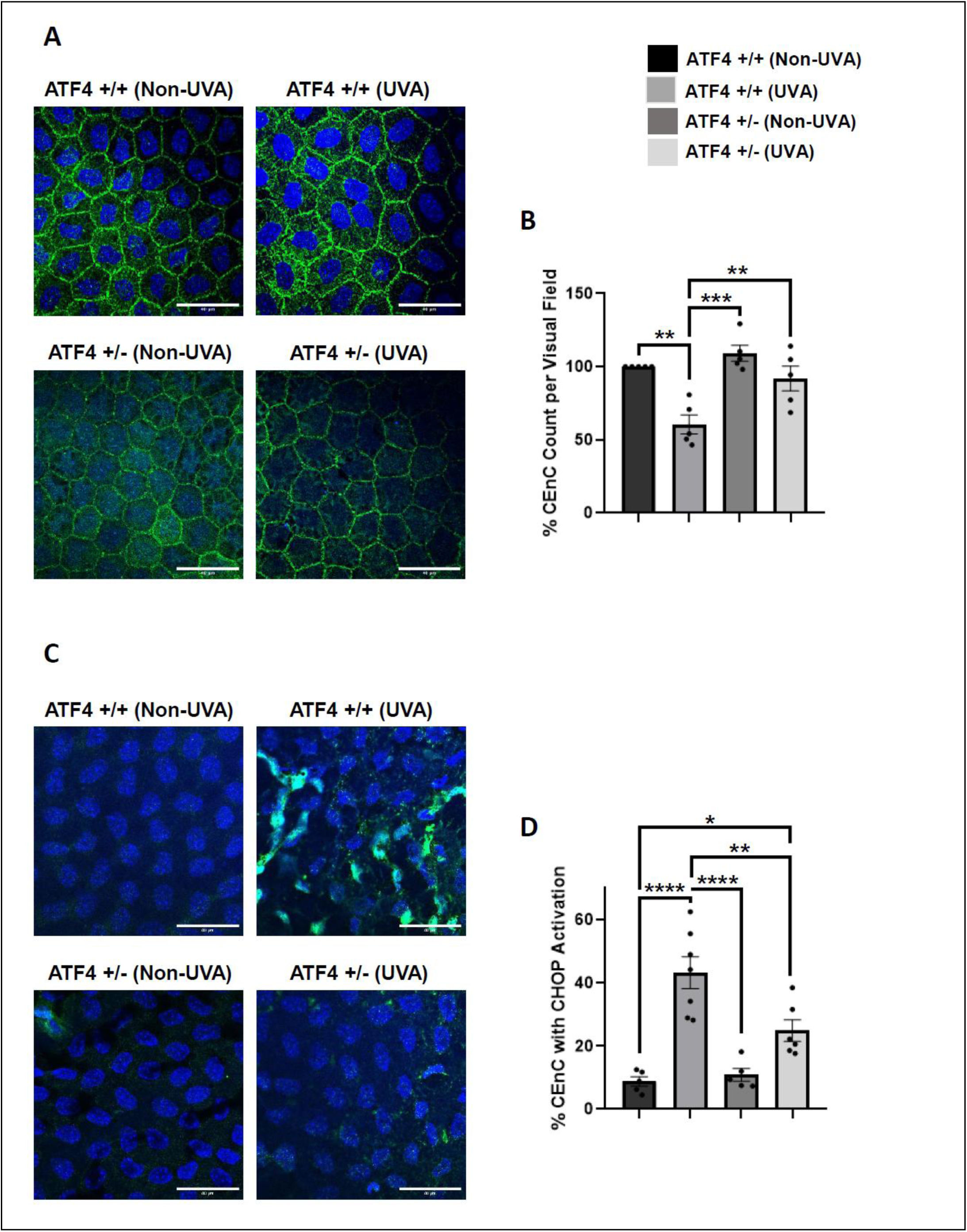
Genetic deletion of ATF4 decreases pro-apoptotic ER stress molecule, CHOP, and protects against UVA-induced cell death. **(A)** Representative Zonula occludens-1 (ZO-1, green) immunostaining showing enlargement and loss of corneal endothelial cells (CEnCs) in *ATF4+/+* compared to *ATF4+/-* mice at week 4 post-UVA (250 J/cm^2^). **(B)** Bar graph representing a significant decrease in CEnC count in *ATF4+/+* compared to *ATF4+/-* mice post-UVA. **(C)** Representative immunostained images for CHOP (green) in *ATF4+/+* and *ATF4+/-* mice CEnCs at day 1 post-UVA. **(D)** Bar graph demonstrating a significant increase in % cells with CHOP in *ATF4+/+* compared to *ATF4+/-* mice post-UVA (n = 5, **P < 0.01, ***P < 0.001, ****P < 0.0001 one-way ANOVA with Tukey’s multiple comparison test) (UVA: 500J/cm^2^, Scale bar: 40 µm).

## DISCUSSION

ER stress and mitochondrial dysfunction have been known to independently contribute to CEnC apoptosis in FECD. Our study reported that tun induces ER stress, which has been supported by numerous studies related to ocular and neurodegenerative diseases.^5,33,34^ In this study, we also investigated the differential induction of pro-apoptotic ER stress and mitochondrial dysfunction in HCEnC-21T and F35T cell lines under unstressed conditions and following tunicamycin. We specifically investigated the role of ATF4 in the PERK-ATF4-CHOP ER stress pathway in mitochondrial dysfunction, mitophagy, and apoptosis in FECD. We demonstrated the upregulation of a well-known ER stress pathway (eIF2α/ATF4/CHOP) in Fuchs’ (F35T) compared to 21T cell lines after Tunicamycin. This is supported by similar findings in many ocular diseases^35–37^, where the higher upregulation of the PERK-ATF4-CHOP ER stress pathway makes cells susceptible to apoptosis. Moreover, in our previous publication, we demonstrated the increasing trend of ATF4 and CHOP in F35T compared to 21T at the baseline. This is supported by similar findings here, where we observe a significant increase in ATF4 and a trend towards increased CHOP levels in F35T compared to 21T under control (DMSO) conditions. A significantly higher level of ATF4 in F35T cell line compared to 21T possibly makes them susceptible to apoptosis, specifically to mitochondrial-mediated apoptosis. This hypothesis is supported by our findings of increased caspase-mediated apoptotic molecules along with mitochondria-mediated apoptosis (Cleaved caspase 9, Bcl-2) in F35T compared to 21T at baseline, which are further upregulated after chronic ER stress. We have not investigated other models of apoptosis, such as ferroptosis under ER or mitochondrial stress in CEnCs, which will be considered in the future.

Activation of mitochondrial-mediated apoptosis is reported to reduce MMP and ATP production and increase mitochondrial fragmentation under pathological conditions. We have observed similar mitochondrial dysfunction where we have increased mitochondrial fragmentation with low ATP and MMP in F35T compared to 21T cell line. Mitochondrial fragmentation and MMP loss are further increased, along with reduced ATP in F35T compared to 21T after tunicamycin. Mitochondrial damage typically triggers mitophagy, which eliminates damaged mitochondria under pathological conditions. PINK1/Parkin-mediated mitophagy is reported in Fuchs’s with activation of Parkin and LC3 and downregulation of PINK1. Our findings support this data, where we first see a significant increase in the expression of Parkin and LC3 in F35T compared to 21T at baseline, suggesting inherent mitochondrial damage in F35T and potentially contributing to its unhealthy state. Additionally, the addition of tunicamycin increased Parkin and LC3 expressions in 21T but not in F35T cell lines. This suggests that the F35T cell line, which is already sick under non-stress conditions, has already activated sufficient mitophagy-initiator molecules. Thus, there is no further effect on mitophagy activation after chronic ER stress, at least in the F35 T cell line. We will explore the effect of chronic ER stress on mitophagy mediators in other Fuchs cell lines in the future. It is also possible that we may need to use either acute ER stress or analyze mitophagy activation at an earlier time point (6 hours), which will help us understand the effect of mitophagy in this F35T Fuchs’ cell line. We will explore the effect of chronic ER stress on mitophagy mediators in less sick Fuchs cell lines with fewer intronic repeats in the future. With respect to PINK1, our data demonstrates significantly decreased expression of PINK1 in the F53T cell line compared to the 21T cell line under non-stress control, as well as in response to Tunicamycin. This finding is consistent with our previous report, which demonstrated downregulation of PINK1 along with upregulation of Parkin at a specific time point after chronic oxidative stress.^38^ In most cases, PINK1 activates Parkin, which tags/ubiquitinates damaged mitochondria for mitophagy. However, there is PINK1-independent activation of Parkin for the initiation of mitophagy,^39^ which could be the case under chronic ER stress and will be investigated in the future.

ATF4 knockdown, either following tunicamycin in vitro or after UVA in vivo, decreased the pro-apoptotic ER stress marker CHOP and reduced cellular apoptosis. These findings are supported by many studies under pathological conditions.^35,40,41^ Our data also demonstrate that ATF4 knockdown reduces MMP loss and mitochondrial fragmentation in 21T after chronic ER stress. Similar studies^16,42,43^ have been reported regarding ATF4 in other cell types under various forms of cellular stress. However, knowledge about the role of ATF4 in mitophagy remains limited.^17^ Even though PINK1/parkin-mediated mitophagy is reported in Fuchs.^38^ In general, it remains unclear whether mitophagy is detrimental or beneficial in FECD. Mitophagy initiates with the recruitment of mitophagy mediators (Parkin, LC3) to damaged mitochondria to form mitophagosomes and culminates with the formation of mitophagic vacuoles containing degraded mitochondria. Mitophagy is considered inhibited if there is a significant accumulation of damaged mitochondria in the form of mitophagosomes.^24,25^ In the late-stage Fuchs disease tissues, a study reports an abundance of autophagosomes with degraded mitochondria, along with activation of PINK1/Parkin^3^, suggesting excessive activation of mitophagy. However, excessive mitophagy has not been confirmed with additional experiments. In our study, although Parkin and LC3 were activated after chronic ER stress in 21T and F35 cell lines, we observed an abundance of mitophagosomes containing damaged mitochondria, suggesting an inhibition of mitophagy. This discrepancy could be because of a specific kind of stress, i.e., ER stress, on a cell line. It is also possible that initial mitophagy mediators (Parkin, LC3) are activating mitophagy. However, extensive mitochondrial damage caused by chronic stress in cell lines probably prevents the completion of all mitophagy steps, resulting in the accumulation of damaged mitochondria in mitophagosomes. We also observed loss of Mfn2 in 21T and F35T cell lines after chronic ER stress. Mfn2 loss is known to inhibit mitophagy, leading to the accumulation of damaged mitochondria as mitophagosomes in cardiac^24^ and skeletal diseases.^25^ There is a possibility that Mfn2 loss is associated with the inhibition of mitophagy in both 21T and F35T cell lines after chronic stress, which may be detrimental to cellular viability. We will explore the role of different stressors in mitophagy for Fuchs in the future.

A previous study revealed that the mitophagy initiator, parkin, is transcriptionally regulated by ATF4.^15^ This finding supports our data, which show that Parkin is activated after Tun treatment under control siRNA conditions. We also reported loss of Parkin after ATF4 knockdown in 21T cell line after tun. Even though Parkin and LC3 are downregulated under ATF4 knockdown after chronic ER stress, suggesting inhibition of mitophagy. We do not observe extensive mitophagosomes, even though the mitophagy initiator Parkin was downregulated under ATF4 knockdown conditions after chronic ER stress, which suggests that mitophagy is very active in this condition. This could be possible due to the rescue of Mfn2 loss under ATF4 knockdown after Tun-induced chronic ER stress, as shown in our data since Mfn2 loss increases mitophagosome formation and inhibits mitophagy in cardiac and skeleton tissues.^24,25^ In the future, we will explore the role of either ATF4 downstream targets, such as CHOP, or other ER stress pathways in mitophagy. Our mouse model of UVA-induced ER stress in FECD demonstrated loss of corneal endothelial cells in ATF4^+/+^ UVA-irradiated mice, while *ATF4^+/-^* mice showed rescue of cell loss post-UVA irradiation. This suggested the role of ATF4 in regulating cell loss *in vivo* as well.

As described above, ER stress and mitochondrial dysfunction have been known to independently contribute to CEnC apoptosis in FECD. However, the role of specific ER stress pathways in regulating mitochondrial dysfunction and mitophagy, as well as their contribution to CEnC apoptosis, remains unknown. Our study provides insight into the crosstalk between ER and mitochondria, as well as the mechanism of ATF4-mediated mitophagy in FECD. One of the limitations of the study is that we need to understand the molecular mechanism by which ATF4 disrupts mitochondria and activates mitophagy. Specifically, it is not clear how ATF4 regulates Parkin to control mitophagy in FECD. It remains unclear whether PINK1 regulates Parkin to control mitophagy in FECD. Moreover, it also remains unclear whether Parkin/PINK-mediated mitophagy pathways are regulated differently under different intracellular stresses. We also need to test the mechanism of mitophagy in detail in animal models of FECD. Other limitations of this study are: 1) F35T is a transformed cell line with upregulated stress by virtue of the model in and of itself as a contributor to some signaling, 2) High-dose UVA mouse model may induce guttae but there is no genetic background of FECD; there is a lack of specificity in this model, particularly because this model’s excessive exposure is associated with damage of other ocular structures in the anterior segment and inflammation that can induce ER stress apart from genetic FECD disease.

In future studies, we will investigate the mechanisms underlying parkin-mediated mitophagy, which contributes to mitochondrial dysfunction and cell loss in FECD. We will also attempt to knock down ATF4 in F35T cells, which will provide more specificity to our model, suggesting ATF4 is involved in FECD. The findings of our study are promising and warrant further testing in FECD animal models. We will also target ER and mitochondrial stress proteins for drug screening and provide therapeutic intervention for ocular diseases such as FECD. In summary, we demonstrate that ATF4 transcription factor plays a key role in ER stress response, which leads to mitochondrial dysfunction and altered mitophagy, thereby causing corneal endothelial cell apoptosis in FECD. By modulating ATF4, we found that ER and mitochondrial stress were alleviated, resulting in enhanced cell survival in the corneal endothelium. This study highlights the role of ATF4 in regulating ER-mitochondrial crosstalk in CEnC degeneration, contributing to FECD development.

## Acknowledgments

We sincerely thank Allison Sowa, Bill Janssen for electron microscopy processing and imaging at The Microscopy CoRE, Icahn School of Medicine at Mount Sinai. We also thank Ula V. Jurkunas (Schepens Eye Research Institute, Harvard University) for providing HCEnC-21T cell line and Albert Jun (Wilmer Eye Institute, John Hopkins University) for providing F35T cell line.

